# Faster and Scalable Parallel External-Memory Construction of Colored Compacted de Bruijn Graphs with Cuttlefish 3

**DOI:** 10.1101/2025.02.02.636161

**Authors:** Jamshed Khan, Laxman Dhulipala, Prashant Pandey, Rob Patro

## Abstract

The exponential growth of genomic data has created an urgent need for scalable sequence analysis algorithms. De Bruijn graphs—along with their colored and compacted variants—have become essential tools in modern bioinformatics pipelines. Colored compacted de Bruijn graphs condense repetitive sequence information, significantly reducing the data burden on downstream analyses like genome assembly, metagenomic clustering, and pan-genomics. Since constructing uncompacted graphs becomes computationally prohibitive at scale, direct construction methods for colored compacted de Bruijn graphs are essential for the scalability of downstream analyses.

We present Cuttlefish 3, a parallel external-memory algorithm that delivers state-of-the-art performance for constructing colored compacted de Bruijn graphs. Our approach introduces three algorithmic innovations that enable efficient scaling to massive datasets while maintaining high performance. First, we develop an optimized technique for accelerating local subgraph contractions. Second, we design a deterministic parallel algorithm based on list-ranking to efficiently merge local solutions. Third, we introduce a novel combinable hash-based method for identifying and tracking color-changing nodes, enabling rapid color-set extraction. We evaluate Cuttlefish 3 on diverse large-scale genomic datasets. In our benchmarks, Cuttlefish 3 achieves 3.29−4.09× speedup over GGCAT, the current state-of-the-art tool, while maintaining comparable memory usage. These performance gains make Cuttlefish 3 a practical solution for representing and analyzing the growing volumes of genomic data in modern bioinformatics workflows. Cuttlefish 3 is implemented in C++17 and is available at https://github.com/COMBINE-lab/cuttlefish.

## 1 Introduction

The de Bruijn graph and its variants are a fundamental representation for analyzing high-throughput genomic data (both raw sequencing reads and assembled genomes). The many uses of the de Bruijn graph throughout bioinformatics are too numerous to enumerate here, but they span a wide gamut, from assembly [1], to variant mapping [2], to sequence-similarity search [3], to indexing [4], to single-cell data processing [5], to pangenome analysis [6].

A primary challenge that sits “upstream” of all of these applications is the construction of the (*colored*) *compacted* de Bruijn graph. The de Bruijn graph is defined on a collection of sequences as a bidirected graph where each distinct k-mer (length-k substring) that appears in some input sequence constitutes a vertex, and each (k+1)-mer forms an edge between its prefix and its suffix k-mers [7,8,1]. The *colored* variant [2] of a de Bruijn graph associates a “color” to each vertex, which is the set of input sequences in which the corresponding k-mer is present in—i.e. the colored variant adds an inverted index to the graph. The *compacted* variant [9] of the de Bruijn graph is obtained by collapsing each maximal non-branching path of the original graph into individual meta-vertices. Likewise, the colored compacted variant associates with each such meta-vertex the relevant color(s). For highly redundant and closely related genomic data, the number of vertices in the compacted graph tends to be *orders of magnitudes smaller* than the original graph [10], which can vastly improve the time- and memory-efficiency of downstream applications.

### Motivation

Yet, the efficiency and scalability gains obtained by using the compacted graph are fully realizable only if we can construct the compacted graph efficiently and directly, without first having to construct the (much larger) uncompacted de Bruijn graph. [9,11,12,13,10,14] for directly constructing these compacted graphs, and the corresponding tools [10] are being used at such a scale, that even moderate improvements in these algorithms can translate into substantial reductions in associated compute costs.

For example, the Logan project [15] has constructed the compacted de Bruijn graph and extracted the constituent unitigs on the entire Sequence Read Archive [16], consisting of >50 PBs of sequencing data (as of the end of 2023) and additionally assembled these into longer contigs. While the tools used for this task are resource-efficient (the Logan project used Cuttlefish 2 [10] for compacted de Bruijn graph construction), the task still required approximately 30 million CPU hours. We estimate that, for the task and type of datasets processed here, Cuttlefish 3 (the new method we propose in this paper) is ≈ 2 times faster than Cuttlefish 2 ^3^, suggesting that this task might instead be completed in closer to 15 million CPU hours. Given that this project used a mix of AWS c6g and c7g instances, costing $0.068 and $0.145 per CPU hour respectively, such computational savings could result in savings on the order of $1.02–$2.18 million, which will only increase with the increasing rate of data production. Similarly, the `Fulgor` [3,17,18] tool provides a state-of-the-art index for colored compacted de Bruijn graphs, and currently uses GGCAT [14] to produce the graph unitigs and color sets upstream for indexing. We estimate that for a range of datasets, close to three quarters of the index construction time is spent in the colored graph construction. Thus a 4 × speedup in this step would lead to about a 2.3 × speedup of the entire construction of the index, a substantial improvement as the indexes grow larger.

### Scalability challenges

Despite significant recent progress on improved algorithms for building colored compacted de Bruijn graphs [9,11,12,13,10,14], existing methods still struggle to scale to extremely large and highly redundant datasets (such as [19]) within feasible timing and resource limits. The primary reason for the scalability challenge in existing tools is algorithmic. Divide-and-conquer based approaches employed by current tools suffer from having to perform a huge number of hash-table queries to traverse subgraphs, which in turn results in sub-optimal performance. Additionally, existing color extraction methods require sorting data volumes proportional to the input size, which further limits the scalability to extreme-scale datasets. In this paper, we propose several algorithmic innovations to address and overcome such issues. The resulting improvements in the graph construction phase would directly improve the efficiency of numerous down-stream applications, e.g., [15,20,3,21,22] among others.

### This paper

In this work, we develop a new tool Cuttlefish 3 for constructing colored compacted de Bruijn graphs that introduces several algorithmic innovations to overcome the challenges in existing tools. Cuttlefish 3 advances the state-of-the-art in this well-studied problem. We demonstrate that it is feasible to scale the construction of colored compacted de Bruijn graphs to very large, multi-terabyte datasets while obtaining high performance and low resource requirements, achieving speedups of up to 4 × over the current state of the art tool, GGCAT [14]. Cuttlefish 3 adopts the “partition-contract-join” paradigm, first introduced over a decade ago [23,9], of partitioning the data into nearly-disjoint subgraphs, performing local contraction with color-set tracking within each of these subgraphs, and then joining together their locally-maximal unitigs into the globally complete graph. While adopting the same overarching paradigm of these approaches, we introduce several algorithmic innovations to demonstrate substantially improved performance and scale to extremely large datasets.

The state-of-the-art performance of Cuttlefish 3 is obtained through novel algorithmic ideas and optimizations, which we believe are of broader interest. Our major algorithmic contributions include: (1) a strategy for reducing the number of graph-membership queries when constructing locally-maximal unitigs within subgraphs; (2) a novel color-set collection scheme by extracting the color information of a very sparse set of vertices using a “combinable hash” of the source identifiers; and (3) a new deterministic and highly parallelizable algorithm for the global stitching phase based on the traditional parallel list-ranking problem [24], inspired from the classical parallel tree-contraction technique [25].

Collectively, the proposed algorithmic techniques yield significant performance improvements for the colored compacted de Bruijn graph construction. For example, our color extraction technique is based on identifying a sparse subset of *color-shifting* vertices (ones which have a different color from their immediate neighbors). By only constructing full color-sets for a subset of the color-shifting vertices, we show that we can obtain the color-sets of all the vertices while drastically reducing the amount of information that must be shuffled and sorted compared to the naive approach of simply collecting and sorting all k-mer–color pairs.

Beyond improving the performance of colored compacted de Bruijn graph construction, the algorithmic techniques we introduce have broader applicability to fundamental problems in parallel and external-memory computing. Notably, we present a novel and practical external-memory solution for the parallel list-ranking problem [24], a classical primitive in parallel algorithms with applications spanning from graph algorithms to computational geometry. While parallel in-memory solutions have existed for decades, the lack of efficient external-memory variants has limited their application to massive datasets. Our deterministic algorithm, inspired by tree-contraction techniques [25], demonstrates that fundamental parallel primitives can be effectively adapted to external-memory settings while maintaining their theoretical guarantees and practical efficiency. Additionally, our combinable hash-based technique for tracking sparse subsets of vertices with differential properties provides a general framework that could be applied to other problems requiring efficient identification and tracking of state changes in massive graphs. We believe that these contributions and ideas extend beyond bioinformatics and could be of independent interest, offering methods for processing large-scale data in domains where memory constraints necessitate external-memory solutions.

## 2 Preliminaries

We assume all symbols to belong to the alphabet Σ = {A, C, G, T }. A *string* s is an ordered sequence of symbols drawn from Σ. |s| denotes the length of s. s_i_ denotes the i’th symbol in s with 0-indexing. The *substring* s_i..j_ of s is the string located within s from its i’th to the j’th indices, exclusive of the index j. The *ℓ*-length *prefix* of s is `pre` _*ℓ*_(s) = s_0..*ℓ*_, and the *ℓ*-length *suffix* of s is `suf` _*ℓ*_(s) =s_|s|−*ℓ*..|s|_. The *append* operation is denoted by · . For two strings x and y such that `suf` _*ℓ*_(x)=`pre` _*ℓ*_(y), the *ℓ-glue* operation ⊙^*ℓ*^ produces the string x⊙^*ℓ*^y=x·y_*ℓ*..|y|_. A k*-mer* is a k-length string.

Each symbol in Σ has a reciprocal *complement*—the complementary pairs being (A, T) and (C, G). For a symbol c, 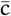 denotes its complement. For a string s, its *reverse-complement* s is the string of the complemented symbols of s in the reverse order, i.e. 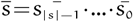. The *canonical* form of s is 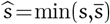, per some strict total order of Σ^|s|^. We lexically order strings.

The *frequency* f(x, 𝒮) of some string x in a multiset 𝒮 of strings is the count of different instances of x present as substrings across all s ∈ 𝒮. Its *canonical frequency* is defined as 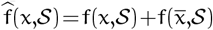

Given a multiset 𝒮 of strings, an integer k > 0, and a frequency threshold f_0_, let 𝒦_+1_ be the set of (k+1)-mers e with f(e, 𝒮)≥ f_0_. The *(edge-centric) directed de Bruijn graph of order* k of 𝒮 is a graph G=(𝒱, ℰ) such that each e = u⊙^k−1^ v ∈ 𝒦_+1_ is an edge from the vertex u to the vertex v. Thus ℰ= 𝒦_+1_ and 𝒱 is induced by ℰ. For ease of notation, we use the terms k-mer and vertex and the terms (k+1)-mer and edge interchangeably.

The *bidirected de Bruijn graph* is an extension of the directed model. ℰ is the set of (k+1)-mers e with 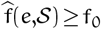. Each vertex represents two k-mers, u and ū, and is denoted with û. It has two *sides*: front and back. An edge e =u⊙ ^*k =1*^ ν is said to *exit* û and *enter* 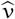. e exits û through its back iff u= û, and it enters 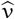 through its front iff 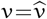. The opposite sides are used respectively in the other cases. Thus the edge e=u⊙^k−1^ ν is represented as, 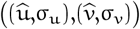 —it goes from the side σ_u_ of vertex û to the side σ _ν_ of vertex 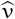. This edge is said to *spell* the (k+1)-me_(_r u⊙^k−1^ ν. The (k+1)-mer 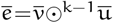 û induces the edge 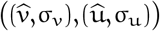 —it mirrors e in the opposite direction [10]. For some side σ, 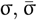 denotes the opposite side.

A *walk* 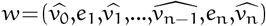 is an alternating sequence of vertexes ν_i_ and edges e_j_ such that any two successive edges are of the form 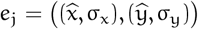 and 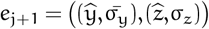. That is, a walk enters and exits a vertex through different sides. Let e_i_ spell the (k+1)-mer s_i_. Then w spells the string s_1_⊙^k^…⊙^k^s_n_. If w is a singleton ^4^, then it spells the k-mer 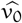. A *path* is a walk w without any repeated vertex, except possibly 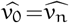, in which case it is also a *cycle*.

A *unitig* is a path 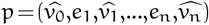 such that either p is a singleton, or: (1) for each 0<i<n, the only edges incident to 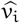 are e_i_ and e_i+1_; and (2) the sides of 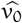 and 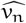 to which e_1_ and e_n_ are incident to, respectively, do not have any other incident edge. A unitig is *maximal* if cannot be extended on either side. The problem of *de Bruijn graph contraction* is to compute all the maximal unitigs of the graph. See [10] for detailed examples of these constructs.

Given the multiset 𝒮 of strings, a *color-set* (or simply, *color*) is a subset of 𝒮. In a *colored de Bruijn graph*, each vertex v has an associated color 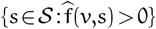—each vertex maps to all the s ∈ 𝒮 in which it is contained. For ease of the algorithm exposition, we redefine colors to be sets of integers: we index 𝒮 as 𝒮 = {s_1_,…,s_n_} and define v’s color as 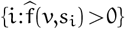

Given some *ℓ* < k and a total ordering of *ℓ*-mers, the *ℓ-minimizer* `mm` (x) of a k-mer x is the smallest *ℓ*-mer substring of x. The *canonical ℓ-minimizer* of x is 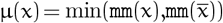. We define `lmm` (x)=µ(`pre` _k−1_(x)) and `rmm` (x)=µ(`suf` _k−1_(x)), the canonical *ℓ*-minimizer of the prefix and the suffix (k−1)-mers, respectively. x is said to be a *discontinuity* k*-mer* if `lmm` (x) ≠ `rmm` (x). A string s with |s| ≥ k is a *super* k*-mer* iff all its substring k-mers have the same canonical *ℓ*-minimizer. ^5^

## 3 Algorithm Overview

We provide a high-level overview of Cuttlefish 3(𝒮,k,f_0_) in this section. The input is a set 𝒮 of strings, an integer k, and a frequency threshold f_0_. The output is the spellings of the maximal unitigs of the induced de Bruijn graph.

A simple algorithm to solve the problem is to first construct the original de Bruijn graph, and then to traverse the graph to find all the maximal non-branching paths. Unfortunately, this approach requires storing a representation of the graph *in-memory*, which quickly becomes infeasible at scale, e.g. for graphs with billions of vertices and edges.

Cuttlefish 3 adopts the partition-contract-join paradigm in which the input data is (1) first *partitioned* into near-disjoint subsets (constituting subgraphs of the de Bruijn graph); (2) all maximal non-branching paths are *contracted* within these subgraphs; (3) and finally the local (per-subgraph) solutions are *joined* into the global solution. These high-level steps have been proposed in this context by [23,9], and later followed by [11,14].

Cuttlefish-3(𝒮,k,f_0_)

// Partition G into a set *G* of subgraphs

1. 𝒢 = Partition-Graph(𝒮,k) // Contract each subgraph 𝒢_i_ and construct the
2. discontinuity graph Γ Γ = Contract-Subgraphs(𝒢,f_0_)
3. // Contract Γ to meta-vertices Π = Contract-Paths(Γ)
4. // Expand back the meta-vertices to Γ and compute the edges’ coordinates χ = Expand-Paths(Γ,Π)
5. // Collate unitigs from the same path in proper rank
6. 𝒫 = Collate-Unitigs(Γ,χ)
7. **return** 𝒫

Cuttlefish 3 first implicitly partitions the graph G into subgraphs by distributing its edges, available from 𝒮, to external-memory buckets using minimizers. Then each subgraph is loaded in-memory and separately contracted using a simple graph-traversal algorithm. The unitig set 𝒰 produced from the subgraphs form partial results of the global problem.

During the local phase, another graph Γ, called the *discontinuity graph*, is constructed in external-memory. The vertices of Γ are the discontinuity k-mers from 𝒮, i.e. it is a subset of the vertex-set of G, and each edge correspond to a unitig in 𝒰. We show that Γ is a collection of disjoint maximal paths and that each one corresponds to a maximal unitig of G. Thus traversing each disjoint path in Γ and collecting the edges in order provides the corresponding global maximal unitig.

At very large scale (e.g., for trillion-base genome collections), using an in-memory representation of Γ also requires substantial space. Instead, we model the problem of computing the “coordinates” of each edge, i.e. the path p it belongs to and its rank in p, as a *list-ranking* problem [26]. We propose a novel algorithm, inspired by seminal parallel algorithms for *tree-contraction* [25], to solve this problem efficiently in external-memory. Once the coordinates of the edges of Γ are available, the full paths, i.e. the global maximal unitigs, are constructed by joining the edges appropriately with a *MapReduce*-like [27] key-value collation scheme, also done in external-memory.

The next subsections describe the steps of the algorithm in more detail. Pseudocodes of different steps and proofs are present in the Supplementary Material. For ease of exposition, we describe the algorithm while skipping more involved details about handling vertex-sides. We emphasize that Cuttlefish 3 and its implementation do take vertex-sides into consideration.

### 3.1 Partitioning G Into Subgraphs

The de Bruijn graph G is first partitioned into a set 𝒢 of 4^*ℓ*^ subgraphs, where *ℓ*<k is the chosen minimizer size, by assigning each induced edge e=u⊙_k−1_ v from the input to the graph 𝒢_rmm(u)_ (which is the same as 𝒢_lmm(v)_). Each edge is thus assigned to a unique subgraph. This partitioning strategy was proposed by the BCALM [11] algorithm. From a given sequence a discontinuity k-mer v participates in two consecutive edges e_1_ =u⊙_k−1_ v and e_2_ =v⊙_k−1_ w, and as e_1_ is assigned to 𝒢_lmm(v)_ and e_2_ is assigned to 𝒢_rmm(v)_, v is assigned in exactly two different subgraphs. Fig. 1a provides an example illustration of such partitioning.

**Fig. 1:**
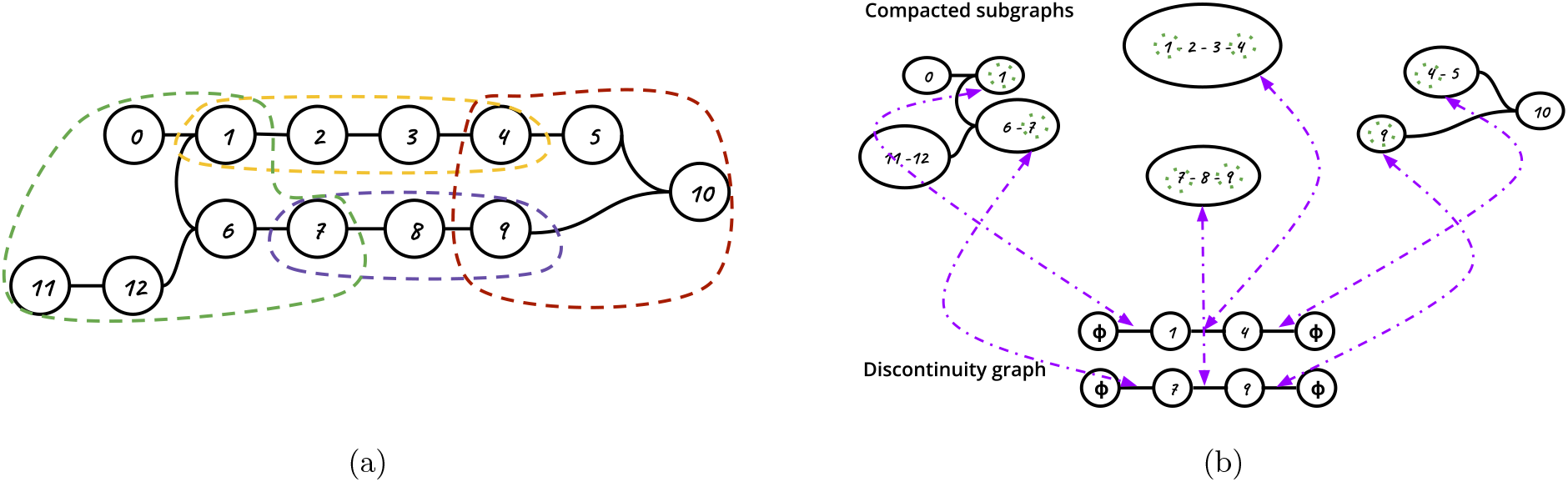
In (a), we show a sample partitioning of a de Bruijn graph G into subgraphs. Each set of vertices enclosed with dashed borders is a subgraph. In (b), we show the compacted subgraphs and the corresponding discontinuity graph (please refer to Sec. 3.2). Maximal unitigs are collapsed into single vertices in the contracted subgraphs. Discontinuity vertices from the original graph are circled with dotted green border. These form the vertex set of the discontinuity graph Γ. Each local maximal unitig containing discontinuity vertex(es) correspond to an edge in Γ, shown with dashed arrows. The maximal paths in Γ correspond to global maximal unitigs of G.

Instead of assigning each edge individually, Cuttlefish 3 assigns longer spans of *consecutive edges*, as induced by their enclosing super k-mers [23], from the input to subgraphs. All the edges from a super k-mer belong to the same subgraph (see Corollary 1). Given a string, all its maximal super k-mers can be computed in linear time using a monotonic queue [28]. We provide a linear time and practically efficient (branch-free) algorithm for such in Sec. 4.1.

### 3.2 Contracting Each Subgraph

Once all input edges are distributed to the subgraphs in external-memory, a hash-table representation of the de Bruijn graph of each subgraph 𝒢_i_ is constructed in-memory. In the representation, the keys are the vertices (k-mers) and their associated values are the *states*—a state is a succinct encoding of the vertex’s neighborhood. Since de Bruijn graphs have bounded-degree, we generalize the vertex-state modeling from Cuttlefish [13] to encode a vertex’s complete adjacency information: frequency-encoding (or presence-absence) of all possible eight incident edges ^6^. For each vertex, Cuttlefish succinctly encodes whether it has unique neighbors at each side and in that case what those neighbors are. Given the state of a vertex, its complete set of edges can be quickly inferred.

Once the in-memory representation of 𝒢_i_ is constructed, the algorithm initiates a non-branching walk from each side of every vertex x. A walk continues so long as, for each extension edge e= {(u,σ_u_),(v,σ_ν_)} from a vertex u, e is the *only* edge incident to the sides σ_u_ and σ_v_. These two walks constitute the maximal unitig of x (and of all the other vertices in the walks). The majority of the work performed during this phase is spent querying the hash table to performing non-branching walks. To extend a walk from u with edge {(u,σ_u_),(ν,σ_v_)}, typical de Bruijn graph traversal strategies [14,29,30,11,31] require probing for the existence of all possible neighbors at σ_u_ and at σν— amounting to eight membership queries for each successful extension of such walks in the DNA alphabet. For unsuccessful extensions, i.e. branching or dead-end at σ_u_ or σ_v_, at least two and up-to eight queries are used. In Cuttle-fish 3, the vertex states in the hash table allows us to elide most of these queries when performing the walks. Specifically, using the neighborhood states of the vertices u and v, Cuttlefish 3 requires *only one query* in a successful extension from u through σ_u_—whether or not σ_u_ or σ_v_ does branch can be inferred from u and v’s states; only v’s state needs to be queried. For unsuccessful extensions, Cuttle-fish 3 requires zero or one query: zero in the case where σ_u_ has no unique edge, and one otherwise, to fetch v’s state. Thus, associating adjacency states to vertices reduces the expected number of membership queries by up to 8 × compared to methods that query for all possible neighbors.

#### Discontinuity Graph Construction

Let x be an end-point of a maximal unitig p in 𝒢_i_ and σ_x_ be the side of x not connected to p. x terminates p due to one of the following reasons: (1) σ_x_ has 0 edges; or (2) σ_x_ has > 1 edges; or (3) σ_x_ connects to a unique neighbor y’s side σ_y_ which has > 1 edges. Maximal unitigs from 𝒢_i_ that have both their endpoints satisfying case (2) or (3) form maximal unitigs of G. Whereas in case (1), if x is a discontinuity vertex, i.e. it has different minimizers i≠ j in its prefix and suffix (k−1)-mers, then the other vertices that would continue the underlying maximal unitig in the global graph G through σ_x_ are absent in 𝒢_i_ and present in 𝒢_j_. 𝒢_i_’s unitigs with endpoints having this property can be further extended in G but not in _i_. Such local maximal unitigs appearing in *different subgraphs* need to be joined together appropriately to compute the correct global output. Cuttlefish 3 solves this problem through another graph Γ, defined from G, called the discontinuity graph. Γ is a weighted and labeled graph with labels associated to each edge. Its vertex set is the set of discontinuity vertices from G. If a maximal unitig p in 𝒢_i_ has discontinuity endpoints x and y, an edge {x,y} is added to Γ with weight 1 and the spelling of p as a label. In case either of x and y is not a discontinuity, a sentinel vertex ϕ is used in its place. Fig. 1b provides an example illustration of contracted subgraphs and the resultant discontinuity graph produced.

#### Construction of Simplitigs

Producing a set of simplitigs [32,33] covering the de Bruijn graph requires trivial updates to this phase of the algorithm. Namely, while processing a subgraph during its compaction, whenever some side of vertex is discovered to be a branching side, the branchingness of it can just be discarded, instead keeping an arbitrarily chosen unique neighbor. Then the subsequent unitig walks can continue along the neighbor through that branching side. This is the same way Cuttlefish 2 produces *maximal path cover* of de Bruijn graphs.

### 3.3 Joining Subgraph-local Maximal Unitigs

#### Discontinuity Graph Structure

We prove that Γ is a collection of maximal disjoint paths (see Lemma 4).

These maximal paths correspond to the maximal unitigs in G that span multiple subgraphs (see Lemma 5).

Consider a maximal path p = (ϕ,v_1_,v_2_,…,v_n_,ϕ) in Γ corresponding to the maximal unitig u in G. Let the sequence of edges in p be (e_1_,e_2_,…,e_n_,e_n+1_). Each e_i_ has a label u_i_ which is a local maximal unitig from some subgraph 𝒢_j_. As such, u=u_1_⊙_k−1_ u_2_⊙_k−1_ …⊙_k−1_ u_n+1_. Hence, if the edges e_i_ of p are known, then ordering the edges e_i_ appropriately and concatenating their labels u_i_ in that order will produce u. But the construction of Γ only produces its edge set {e:({x,y},1,s)} (see Sec. 3.2), and hence for some e_i_, we know neither the ID of the maximal path p it belongs to ^7^, nor its rank i in p. We model this problem in detail as following.

#### Formulating a List-Ranking Problem

Let p=(ϕ,v_1_,v_2_,…,v_n_,ϕ) be a maximal path in Γ. For simpler exposition, let us differentiate its two instances of the sentinel vertex ϕ as p=(v_0_=ϕ_l_,v_1_,v_2_,…,v_n_,v_n+1_=ϕ_r_) ^8^. Let p’s edges in order be (e_1_,e_2_,…,e_n_,e_n+1_). For each e_i_, we need to compute the ID of this path p and its rank in p. The (0-based) rank of e_i_ in p’s edges is `rank` (e_i_)=i−1, whereas the rank of v_i_ in p’s vertices is `rank` (v_i_)=i. Since e_i_ =({v_i−1_,v_i_},,), `rank` (e_i_) =`rank` (v_i−1_). Thus we can solve for the vertices’ ranks instead. Considering each maximal path p an ordered list, we need to compute for each vertex v, which list (path) it belongs to, `path` (v), and at what rank, `rank` (v)—this is exactly the classical *list-ranking* problem [26].

#### Solving the List-Ranking Problem

There exist parallel in-memory solutions [24] for the list-ranking problem. But with practical datasets at scale, this graph can become very large for an in-memory representation. Therefore, we design a parallel and external-memory solution for this problem instance, inspired from the algorithm design technique of *tree-contraction* [25]. The idea is to (in parallel) contract the list to a single vertex through a sequence of *compress* operations, which essentially contract sets of vertices into single vertices, and solve the problem of interest incrementally during contractions.

We aim to contract each maximal path p of Γ into a single vertex v through a series of contractions and solve the list-ranking problem for just v. Then, we aim to expand v back to p through a series of expansion operations, in the opposite order of the contraction operations, and solve the problem for the rest of the vertices of p along the way. We will perform these contraction and expansion operations in batches, keeping only a disjoint subset of the vertices and the edges of Γ in memory for each batch, thus avoiding a full in-memory graph representation.

#### Discontinuity Graph Contraction

We hypothetically partition the vertex set of Γ into n partitions—each vertex v belongs to a partition 𝔓 (v) ∈ [1,n] through uniform hashing, with 𝔓 (ϕ_l_)= 𝔓 (ϕ_r_)=0. We load each partition 𝒫_i_ into memory, sequentially, in descending order from n to 1 (though any order would work) and contract each ν ∈ 𝒫 _i_ and their incident edges from Γ in parallel, as follows.

Suppose that a maximal path p has the form (…,x,y,z,…) right before contracting partition 𝒫_i_. Then 𝔓 (v) ≤ i for all v ∈ p, since all the partitions 𝒫_j>i_ have been contracted by this point. Say that y belongs to 𝒫_i_ but not x and z, i.e. 𝔓 (y) =i, 𝔓 (x) <i and 𝔓 (z) <i, and the edges between them are e_xy_ =({x,y},w_xy_,) and e_yz_ = ({y,z},w_yz_,). Then we merge the edges e_xy_ and e_yz_ to a new edge e_xz_ =({x,z},w_xy_+w_yz_,). If there are successive vertices (y_j_) in p = (…,x,y_1_,…,y_m_,z,…) with 𝔓 (y_j_)=i, then the entire chain of y_j_ is merged to a single edge e_xz_ having weight equal to the sum of the weights along the chain’s edges. This way all the edges incident to 𝒫_i_ are contracted in parallel and a smaller list is produced for the next step of contracting 𝒫_i−1_.

Finally, during contraction of 𝒫_i_, if some path p is found to be of the form (ϕ_l_,y,ϕ_r_), we set y to be the path-ID of p. Hence, `path` (y)=y and `rank` (y)=w_ϕ ly_, the weight of the {ϕ_l_,y} edge. Since compacted de Bruijn graphs form a vertex-decomposition of the graph [11], y is uniquely present in this contracted path (and the original maximal path)—the strategy of assigning each maximal path in Γ the ID of one of its vertices as its list ID ensures uniqueness of the list IDs. The rank of y is correct by induction as all edges in the original path have unit weights and the contractions merge edges by summing their weights. Fig. 2a illustrates an example execution of this method.

**Fig. 2:**
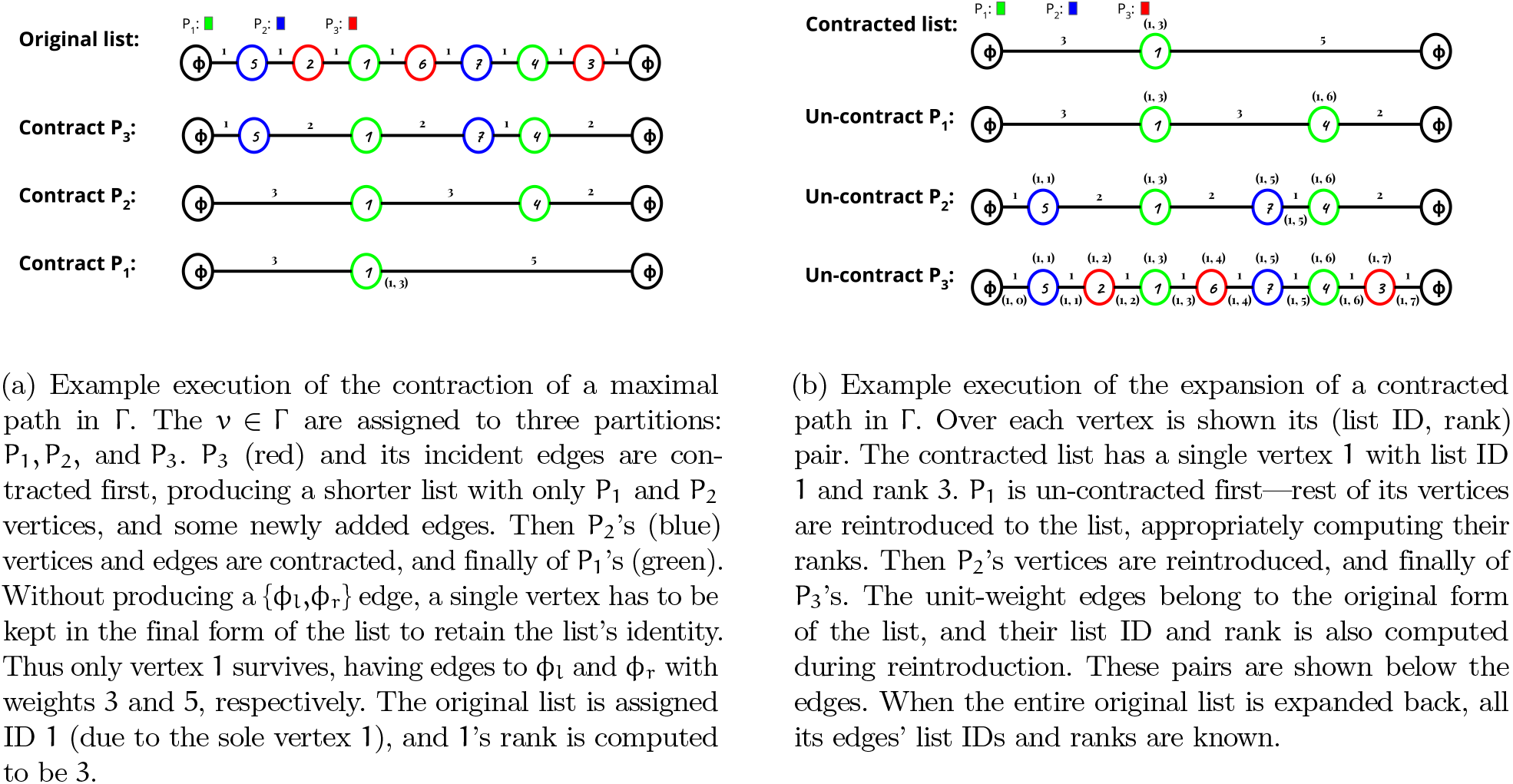
Example contraction and expansion of a maximal path in Γ.

#### Expansion of the Contracted Graph

Now that for each maximal path, the path ID and the rank of a single vertex is computed, we use this partial solution incrementally to compute the solution for the rest of the vertices. We use an expansion (un-contraction) scheme that is complementary to the contraction process to expand back the contracted paths to their original forms. The algorithm loads each partition 𝒫_i_ into memory sequentially in the opposite order that the partitions were contracted, i.e. in ascending order from 1 to n. By induction, the path ID and the rank of each vertex in 𝒫_i_ have been computed at this point.^9^ The base case of the induction is for 𝒫_1_—all its vertices’ path IDs and ranks have been computed in the last step of contraction. Now during the i’th step of expansion, for each v ∈ 𝒫_i_, `path` (v) and `rank` (v) are propagated through its incident edges, including the ones introduced during the contractions, in parallel, as follows.

Say that for vertices x, y, and z, there exist edges e_xy_ =({x,y},w_xy_,) and e_yz_ =({y,z},w_yz_,). These edges may belong to the original path, or may have been introduced by the contractions. Suppose that 𝔓 (y) = i, 𝔓 (x) > i, and 𝔓 (z) > i. Since y belongs to _i_, its path ID and rank have been computed by now. Let these be `path` (y) =π and `rank` (y) =ρ, respectively. Then during expansion from _i_, e_xy_ and e_yz_ are reintroduced to some path p. The path ID and the ranks of x and z are computed as `path` (x) =`path` (z) =π, `rank` (x) =ρ−w_xy_, and `rank` (z)=ρ+w_yz_. Besides, if any of e_xy_ and e_yz_ has weight 1, then it is an original edge of Γ and we need its path ID and rank. As described in the list-ranking formulation, for such an edge e = ({u,v},1,s), `path` (e) = `path` (u) and `rank` (e)=`rank` (u). The correctness of the computed path IDs and the ranks during expansion hold by induction. Fig. 2b illustrates an example execution of the method.

#### Discontinuity Graph Representation

For the proposed path contraction and expansion strategies to work, we require the ability to load partitions of Γ into memory. Recall that, during subgraphs contraction (see Sec. 3.2), Γ is available as an edge-list. Edge-lists, as is, do not support loading edges only incident to a specific vertex partition without a complete scan. As such, we use a modified variant of the edge-list to represent Γ, which we refer as the *blocked edge-matrix*.

The blocked edge-matrix of Γ is an (n+ 1) × (n+ 1) matrix where each cell is an edge-list, which is a block (subset) of the actual edge-list. Each cell has an associated external-memory bucket for its edges. Consider an original edge e = ({x,y},1,s) in Γ, and let 𝔓 (x) = p and 𝔓 (y) = q. We add e to the (p,q) and the (q,p)’th cells of Γ. Newly introduced edges during contraction of Γ are also added to the matrix similarly. Thus when we need to access all the edges incident to partition i, we only load the edge-lists in either the i’th row or the i’th column of Γ.

We further reduce the external-memory requirement of this representation by half by adding e to only one cell. For e=({x,y},w_xy_,), let p= 𝔓 (x) and q= 𝔓 (y). WLOG, assume p≤q, otherwise we swap the endpoints. We add e only to the cell (p, ≤ q).

Recall that, during contraction, when processing 𝒫_j_, we only need access to the edges between (𝒫_i_, 𝒫_j_) for all i≤ j. Edges between (𝒫_j_, 𝒫_k_) for k>j are not required as 𝒫_k_ has been contracted out by this point. Then it suffices to only load the cells (i,j) such that i≤j—which are in the j’th column of Γ in its upper triangle. Similarly, during expansion, when processing 𝒫_i_, we require access to only the edges between (𝒫_i_, 𝒫_j_) for all j ≥ i. Edges between (𝒫_h_, 𝒫_i_) for h<i are not required as 𝒫_h_ has already been reintroduced and solved by this point. As such, loading only the i’th row of Γ in its upper triangle is sufficient.

Since at each iteration of the contraction and expansion procedure, we only need to stream through some row or column of this matrix, this algorithm is completely amenable to the external-memory setting. Fig. 3 provides an example state of the blocked edge-matrix and the contraction of a partition.

**Fig. 3:**
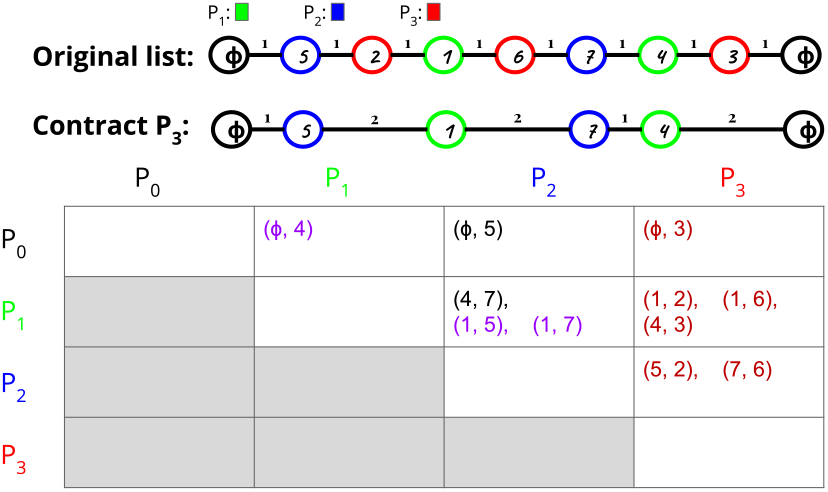
Upper-triangular edge-matrix for an example discontinuity graph Γ with only one maximal path (list), and a single contraction step. Red entries correspond to edges incident to P_3_. These are “merged” when contracting P_3_, resulting into the purple entries introduced to Γ.

#### Collation of the Local Maximal Unitigs

After the expansion of contracted Γ, the path ID and the rank of each original edge of Γ have been computed. These pieces of information reside in different cells of the blocked edge-matrix. For an original path p=(v_0_,e_1_,v_1_,…,v_n_,e_n+1_,v_n+1_), e_i_’s information is in the cell (𝔓 (v_i−1_), 𝔓 (v_i_)) of Γ (or in the transpose cell). The e_i_’s need to be collated together from the different cells in their proper rank, and their labels then concatenated, to produce the corresponding maximal unitig of p. We perform this with a *MapReduce*-like [27] scheme. For each original edge e = (,1,s) with `path` (e) = π, `rank` (e) = ρ, and label s, we consider π as its key and (ρ,s) as its value. Edges with the same key are *mapped* to the same bucket, and each bucket is then *reduced* to bring all the edges with the same key in it together in their appropriate rank.

We allocate a list KV of η external-memory buckets for the original edges. For each original edge e=(,1,s) in parallel, we map e’s path information (π,ρ) and its label s to the bucket KV^ξ(π)^, where ξ: Σ^k^ →[1,η] is a uniform hash function. Then for each bucket KV_i_ in parallel, we reduce it: we sort KV_i_ by the key (π,ρ), which brings together all the labels with the same π-value in their ρ-order. Concatenating the labels s with the same π-value in this order produces the maximal unitigs of G that span multiple subgraphs.

### 3.4 Colors Extraction

The colored de Bruijn graph adds an inverted index structure to the graph: each vertex is associated to its *color* and thus maps back to all the input sources in which it is contained. To extract the colors of all the vertices, the scheme used in existing methods [14,29,34] is to collect all the pairs (v,s) such that the vertex (i.e. k-mer) v appears in source s, then to sort these pairs to collate the sources of each vertex. Each k-mer instance from the input constitutes a pair in this approach. At scale, this entails sorting substantial amounts of data, either as a whole [29,34], or in parts [14,35].

In Cuttlefish 3, we introduce a novel method for collecting the unique colors of a graph and associating the vertices to these sets. We avoid collecting the color information for the entire vertex set, and instead propose a scheme to collect the colors for only a *sparse* subset of the vertices, from which the colors of the remaining vertices can be directly inferred based on the contracted form of each subgraph 𝒢_i_. During the construction and contraction of 𝒢_i_ (see Sec. 3.2), a subset ϒ_i_ of its vertices are identified as of interest. These are the vertices where the maximal unitig walks encounter shifts in colors ^10^: for each subpath p with the same color c in a maximal unitig, only the first vertex v is added to ϒ_i_. If c can be extracted for v, then the color of the entirety of p is available. However, this forms a causality dilemma—we need to identify the shifts in colors in 𝒢_i_’s maximal unitigs in order to construct ϒ_i_, so that 𝒢_i_’s vertices’ colors can be collected.

To address this, we use the fact that we only need to detect the *shifts* in colors in a maximal unitig to construct ϒ_i_, and not the colors themselves. For a color c={j_1_,…,j_n_, we define its *signature* as h(c) = h^*′′*^ h^*′*^(j_1_),…,h^*′*^(j_n_) h^*′*^ is a hash function for the source-ID j’s and h^*′′*^ is an *online* ^11^ hash function to combine h^*′*^’s values.

During the construction of 𝒢_i_, h^*′′*^ being online enables us to compute h(c) for each vertex v in 𝒢_i_, without having its full color c present. During partitioning (Sec. 3.1) we associate to each edge e the source ID j from which it originated. Then while constructing 𝒢_i_ (Sec. 3.2), for each edge-source pair (e={u,v},j), the color signatures associated to the vertices u and v are updated online with h^*′*^(j). ^12^

During the contraction of 𝒢_i_, as each maximal unitig p is constructed, a walk over p provides the vertices where h(c) shifts. The set of unique colors across the whole graph G is tracked with a concurrent hashtable, 𝒞 which maps a color signature h(c) to the coordinate (or pointer) of c in the color-output. For a color-shifting vertex v in p, if its h(c) is not found in 𝒞, then it is added to ϒ_i_ as a vertex of interest. Otherwise, c’s coordinate in the color-output (available from 𝒞) is associated to v’s position in p.

After the contraction of 𝒢_i_, all vertices in 𝒢_i_ that are color-shifting *and* have their colors absent in C are identified and present in ϒ_i_. Then we scan the edge-source pairs (e= {v_0_,v_1_},j) of 𝒢_i_ again, and for each v found to be in ϒ_i_, we add the key-value pair (v,j) to an external-memory collection. It is then semi-sorted with a *counting sort*-like method to group together pairs with the same keys—the values in a group with the key v constitute v’s color c. c is added to the color-output, and the key h(c) is added to with 𝒞 c’s coordinate in the output as its value. We also associate this color-coordinate back to v’s position in its maximal unitig p (these positions are tracked separately).

Finally, when the locally maximal unitigs are collated (Sec. 3.3), their colors are also gathered and collapsed if required—consecutive unitigs can continue sub-paths with the same color. Fig. 4 illustrates an example of the color signatures and how these are computed online.

**Fig. 4:**
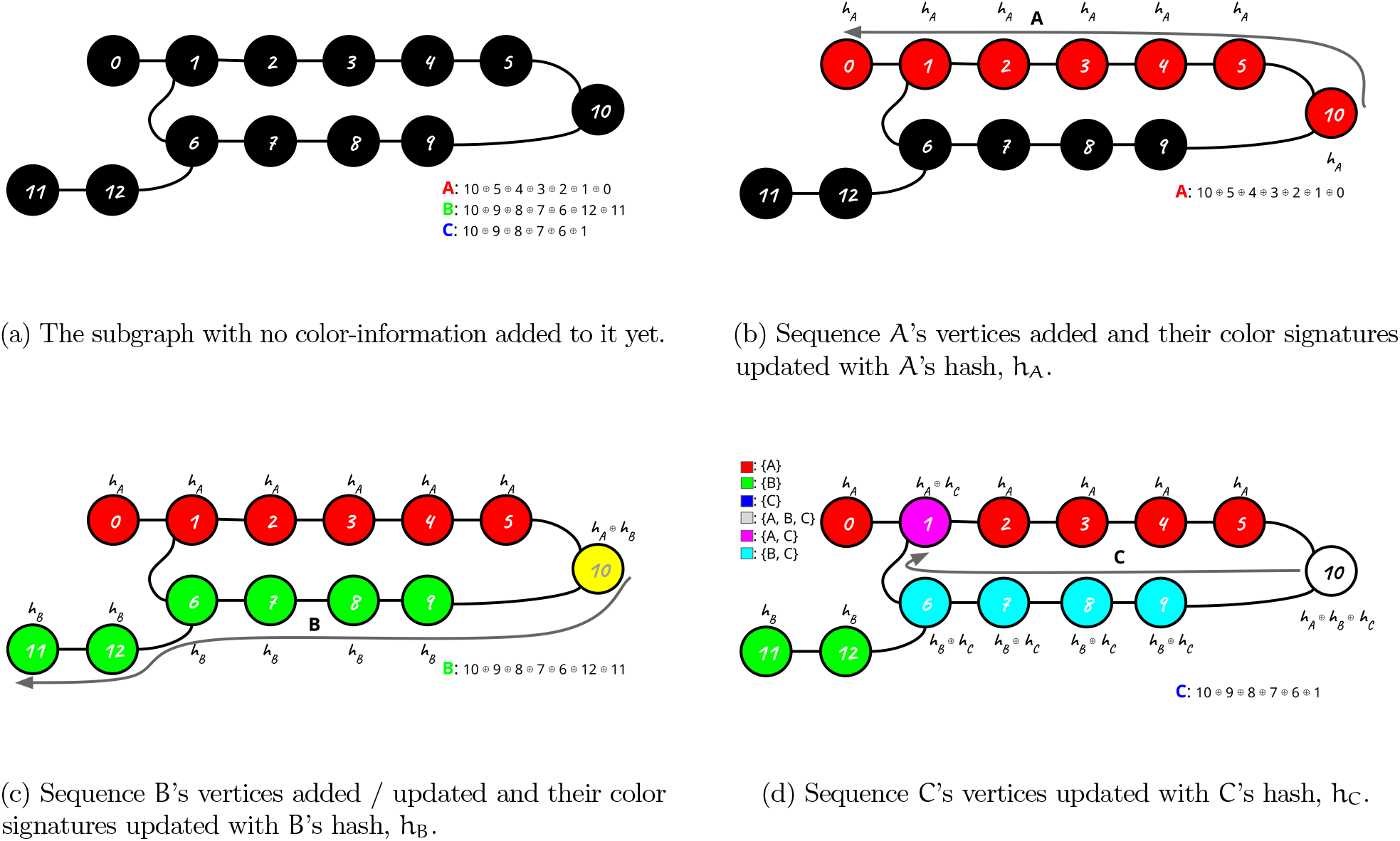
A subgraph constructed from three sequences A (red), B (green), and C (blue). The sequences are denoted as concatenations of vertices (k-mers). The figures show how, during the construction of the subgraph, its vertices’ color signatures (color-coded visually) get updated. The vertices have no color-information at the beginning, color-coded as black. Then the sequences A, B, and C are traversed in order. With each vertex seen during the walks of the sequences, its color signature is updated online. Finally, the complete color signature is available for each vertex. The color-shifting vertices are the ones where either the unitig walks begin (so a new color-stretch starts) or where the walk encounters a new color: 0, 1, 2, 6, 10, and 11.

#### Asymptotics

We briefly discuss the running times of the different phases of the algorithm. For an input set 𝒮, let its total size be *ℓ* = Σ_s∈ 𝒮_|s|, the number of vertices (unique k-mers) and edges (unique (k+1)-mers) in the de Bruijn graph G be n and m respectively, and the number of vertices in the discontinuity graph Γ be γ. The partitioning step makes one pass over each s∈ 𝒮, touching each k-mer instance once, requiring 𝒪 (*ℓ*) work. Next, constructing a subgraph 𝒢_i_ requires scanning over 𝒢_i_’s edges (or groups of edges through super k-mers), as many times as they are present in the input—in total amounting to another scan over the input, 𝒪 (*ℓ*). Then the contraction of 𝒢_i_ traverses it once, requiring 𝒪 (n+m) work in total. Then we process (contract and then expand) Γ, where each vertex and its incident edges are processed twice—once during contraction and once during expansion. Also, each maximal path p in Γ produces (|p|−1) new edges during contraction, summing to 𝒪 (γ) new edges. In total, these steps require work 𝒪 (γ). Collation of the locally maximal unitigs involve moving them to appropriate buckets and to semi-sort them. Since they sum to (n+γ) in size, this step requires 𝒪(n) work.

## 4 Algorithm Optimization

We discuss some of the salient optimizations and implementation details that have been critical to optimize Cuttlefish 3’s performance in this section.

### 4.1 Branch-Free Minimizer Computation

Instead of distributing the edges to subgraphs piecewise, we group together consecutive edges in the input as maximal super k-mers [23], similar to [14,11]. Given a string s, splitting it into maximal super k-mers requires computing the minimizer of each constituent k-mer. Computing the *ℓ*-minimizer of a k-mer takes time 𝒪 (k−*ℓ*). Naively computing all minimizers in s takes time 𝒪 (|s|(k−*ℓ*) . This is typically improved to amortized time 𝒪 (|s|) as following. The k-mers of s are scanned, while a deque q of *ℓ*-mers is maintained. q stores a subset of the *ℓ*-mers of the current k-mer, that are *candidate* minimizers for k-mers yet to be scanned. Each subsequent k-mer v introduces a new *ℓ*-mer x to q, while the oldest *ℓ*-mer z in q, if out of range of x, is removed from q. q is maintained sorted—all *ℓ*-mers y>x in q are popped off as they cannot be the minimizer of v or any next k-mer. The front of q thus contains the minimizer of v. Since each new *ℓ*-mer x is compared with a variable number of y ∈ q and the resulting minimizer is available only after such comparisons, this method can induce many *branches* in the generated processor instructions.

Instead, we simply maintain an ordered list M of all the *ℓ*-mers of the current k-mer, and compute the minimum of M for each k-mer. Computing the minimum of M from scratch takes time 𝒪 (k−l), but we reuse the computations for subsequent k-mers. A k-mer has (k−*ℓ*+1) *ℓ*-mers, and two consecutive k-mers u and v share (k−*ℓ*) *ℓ*-mers. Say the list of the last (k−*ℓ*) *ℓ*-mers of u is M_u_. v has just one new *ℓ*-mer y apart from M_u_. If we know x = min(M_u_), then v’s minimizer computation is just min(x, y)—a single computation doable without branches.

We partition M into two windows: L and R. L is at the front of M while R is at the back. Initially, L=M and R is empty. So min(M)=min(L). For each new k-mer v, an *ℓ*-mer y is appended to R, while one *ℓ*-mer falls out of L’s prefix. So a suffix L_suf_ of the initial L and the entire current R belong to v. We construct a suffix-min table for the initial L, computed at the beginning, from which min(L_suf_) is available. We also maintain the running min(R), updated with every new addition to it. Thus the minimizer of v is min min(L_suf_),min(R) . This computation is branch-free.

When L becomes empty, we reset the data structures: L and R are swapped ^13^, and we recompute the suffix-min table of L. This computation has one linear scan computing a running minimum, which introduces some conditional dependence among instructions. This can be minimized with vectorization.^14^.

The suffix-min table computation takes time 𝒪 (k−l+1), happening at every (k−l+1)’th k-mer. Thus the total time to compute all the minimizers is 𝒪 (|s|), and each takes amortized constant time.

### 4.2 Cache-friendly Subgraph Atlases / B-tree

With a separate subgraph for each *ℓ*-minimizer, we have to maintain 4^*ℓ*^ subgraphs. With practical values of *ℓ*, handling many external-memory buckets poses bottlenecks with the filesystem. So we group together minimizers into a smaller number of subgraphs through uniform hashing.

It is also preferable to have a large enough number g of subgraphs such that the subgraph sizes are small enough to avoid large in-memory representations. In a parallel implementation with w workers, each subgraph bucket should have w worker-local buffers of super k-mers so as to minimize lock-contention for the subgraph. This results in g × w buffers in-memory, and the end of each are contending for space in cache. To improve caching behavior, we group together subgraphs into *atlases* (similar to texture-atlases [39]). We keep 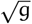 atlases, each containing 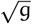 subgraphs, resulting into our g original subgraphs. Super k-mers are added to the atlases, instead of directly to the subgraphs, so that we maintain only 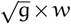 buffers in memory. Only when a buffer fills up are its super k-mers distributed to the 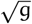 subgraphs in that atlas. In that case, an additional 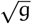 buffers’ ends come into the cache. Note that this scheme is similar to having a *B-tree* of the subgraphs. A small example figure demonstrating this idea is provided in Figure S1.

### 4.3 Color-signatures for Multisets

Defined in Section 3.4, the signature of a color c is h(c) = (h^*′′*^ h^*′*^(j_1_),…,h^*′*^(j_n_))—it is a hash-combination of the hash values of j ∈ c. A vertex v can belong to the same source s_j_ multiple times, and as such, its color c will be observed in a multiset variant in this case. Thus h^*′*^(j) will be combined to h(c) multiple times. h^*′′*^ being online and memoryless, the color c can have different signatures for the different variants of c. This makes it impossible to correctly identify the color-shifts.

To tackle this, we process the input sources in a consistent order, so that the subgraph buckets contain their super k-mers in that order. As a result, any multiset variant of c is always observed in sorted order. Each vertex ν in 𝒢_i_ keeps track of the last source ID j added to its h(c), and if j is added again to it, we simply skip it. This ensures that all multiset variants of c produce the same h(c).

## 5 Results

We evaluate the performance characteristics of Cuttle-fish 3 ^15^ and how it compares to the state-of-the-art solution GGCAT [14] ^16^ in constructing (colored) compacted de Bruijn graphs in this section. We validate the correctness of the output produced by Cuttlefish 3 by comparing it against naively constructed solutions for multiple datasets. In the naive construction, we first build the de Bruijn graph in-memory and then compact it by traversing the graph.

Experiments were performed on a single machine with four Intel(R) Xeon(R) Platinum 8160 2.10GHz CPUs having 96 cores in total, 1.5 TBs of 2.66 GHz DDR4 RAM, and two 3.5 TBs Toshiba PX05SRB384Y ATA SSDs. The running times and the maximum memory usages were measured with the GNU `time` command. The evaluated tools were restricted to CPU-usages concording to the number of threads with the `cpulimit` tool^17^.

### 5.1 Datasets

The experiments are performed on the following datasets: (1) Human gut [40]: 30,691 representative sequences from prevalent human gut prokaryotic genomes (≈ 61B bp, 18 GB gzipped FASTA); (2) Salmonella (309K): 309,123 *Salmonella* genome sequences used in [14], downloaded from the *EnteroBase* database (≈ 1.5T bp, 1.36 TB FASTA); and (3) Bacterial archive [19]: 661,405 bacterial genomes (≈ 2.58T bp 757 GB gzipped FASTA).

### 5.2 Colored Compacted de Bruijn Graph Construction

We evaluate Cuttlefish 3 compared to GGCAT with k = 31 to construct the colored de Bruijn graph on the genome sequence datasets. [14] demonstrates that GGCAT is more efficient than Bifrost for this use-case and as such we do not compare against it. Table 1 contains the benchmark results. Supplementary table S1 additionally contains the disk-usage results.

**Table 1.** Time (in wall clock) and memory (in GB, in parentheses) performance results for constructing colored compacted de Bruijn graphs with 32 threads. The best metrics in each row are highlighted.

| k | Dataset | GGCAT | CUTTLEFISH 3 |
| --- | --- | --- | --- |
| 31 | Human gut | 19 m (**20.6**) | **11 m** (25.2) |
|  | Salmonella (150K) | 02 h 18 m (10.4) | **42 m** (**10.3**) |
|  | Salmonella (309K) | 05 h 18 m (**14.0**) | **01 h 32 m** (15.9) |
|  | Bacterial archive (661K) | 13 h 29 m (35.8) | **03 h 18 m** (**33.7**) |
| 63 | Human gut | 15 m (22.2) | **11 m** (**20.4**) |
|  | Salmonella (150K) | 02 h 21 m (**10.1**) | **39 m** (11) |
|  | Salmonella (309K) | 05 h 39 m (**14.2**) | **01 h 30 m** (17.5) |
|  | Bacterial archive (661K) | 11 h 25 m (**36.2**) | **03 h 15 m** (41.3) |

From Table 1, we note that Cuttlefish 3 is significantly faster than GGCAT for all the datasets. On 150K Salmonella genomes, Cuttlefish 3 is ≈3.29–3.62× faster than GGCAT. For 309K Salmonella, Cuttlefish 3 is ≈3.46–3.77 ×faster. On the bacterial archive (661K), it is ≈3.51–4.09 × faster than GGCAT. The memory usages are roughly similar across the test configurations.

### 5.3 Parallel Scaling

We assess the parallel scaling speedup of Cuttlefish 3 across a varying number of processor-threads in this experiment. We use a subset of 10,000 bacteria from the bacterial archive dataset for this. We set k=31, and executed Cuttlefish 3 with thread-counts ranging in 1–32. Figure 5 demonstrates the running times. We also include in this figure an estimated loose lower bound on the minimal time achievable for construction on this dataset at different thread counts. This estimate is obtained by simply measuring the time required to (in parallel) decompress the input files twice, but without performing any parsing. It seems reasonable that any approach—including those using the partition-contract-join paradigm—will have to scan the data at least two times, motivating our proxy measurement for a loose lower bound.

**Fig. 5:**
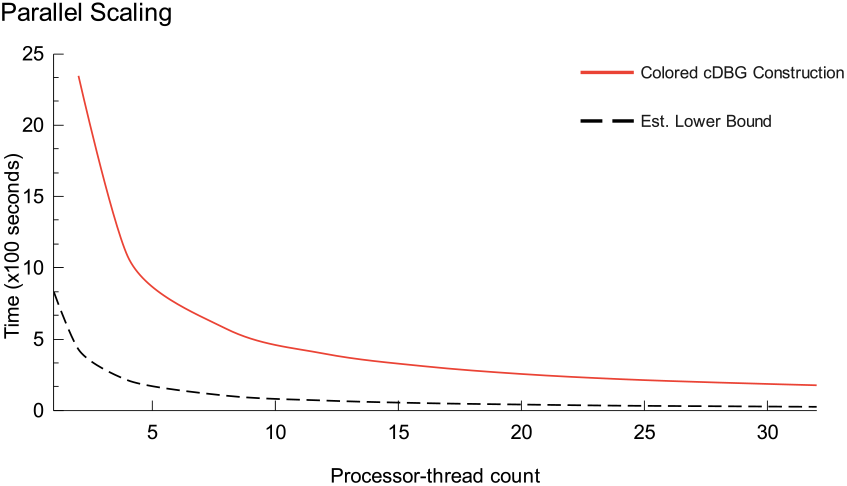
Time incurred by Cuttlefish 3 across varying processor-thread counts for 10K bacteria.

Supplementary Table S2 provides parallel scaling performance of Cuttlefish 3 compared against GGCAT. The GGCAT implementation did not work successfully in our benchmarks for thread-counts that are not powers of two, so we provide performance statistics for powers of two, up to 32..

### 5.4 Sparsification for Color Extraction

In the colored compacted graph construction experiments, we measured the effectiveness of our sparsification strategy (outlined in Section 3.4). Our strategy not only reduces the number of vertices for which the full color needs to be extracted but also exploits the fact that many color-shifting vertices will have identical colors. If a color corresponding to the signature (i.e. hash) of a color-shifting vertex has already been computed and inserted into the color-table, then it is unnecessary to assemble that color again. Table 2 provides some statistics on the degree of sparsification, and thus reduction in overall work, that our strategy enables. We observe that the number of color shifting vertices is usually a small fraction of the total number of vertices (i.e. the graph size). More importantly, even for the vast majority of color shifting vertices, at the time we encounter them, we have already computed the color corresponding to their signature. Thus, across the three datasets in Table 2, we need to compute the colors for only between 0.83%−3.78% of the total vertices in the graph.

**Table 2.** Graph and color related statistics for the datasets for which we construct the ccdBG.

| Dataset | Graph-size<br>($\times 10^9$) | Color-shifting<br>vertex<br>($\times 10^9$) | Vertices w/<br>new color-<br>signature<br>($\times 10^9$) | Percent-<br>age | Pairs<br>sorted<br>($\times 10^9$) | Colors<br>($\times 10^6$) |
| --- | --- | --- | --- | --- | --- | --- |
| Human<br>gut (31K) | 15.33 | 1.95 | 0.232 | 1.51 | 10.05 | 227.8 |
| Salmonella<br>(150K) | 0.54 | 0.08 | 0.019 | 3.63 | 478.0 | 19.3 |
| Salmonella<br>(309K) | 0.72 | 0.109 | 0.027 | 3.78 | 1109.3 | 27.2 |
| Bacteria<br>(661K) | 48.78 | 5.71 | 0.404 | 0.83 | 1459.8 | 389.1 |

## 6 Conclusion

We have introduced Cuttlefish 3, a fast, parallel, and low-memory algorithm, and its prototype implementation to construct colored compacted de Bruijn graphs. It is highly scalable across a variety of large-scale genomic datasets. We observed that Cuttlefish 3 constructed the colored compacted graph for massive datasets up-to 4.09 × faster than the current state-of-the-art method, GGCAT, with the same memory profile. Cuttlefish 3 reduces Cuttlefish 2’s computational requirements substantially by adopting the partition-contract-join paradigm initially introduced by BCALM 2. Moreover, it introduces several novel ideas to this paradigm. Cuttlefish 3 generalizes the vertex-modeling schemes of Cuttlefish 1 and 2 to associate neighborhood states to vertices in subgraphs, which significantly reduces the amount of work to traverse the subgraphs. It makes use of a novel algorithm to stitch together the local solutions from the subgraphs. For colored graphs, it introduces a novel multi-pass scheme to vastly sparsify the set of vertices only for which colors need to actually be tracked and constructed, which is sufficient for the entire compacted graph. For example, we found that while tracking the colors for only 0.83% of the unique k-mers of the 661K bacterial archive [19], Cuttlefish 3 is able to construct the correct colors for all of the vertices.

In addition to the algorithmic improvements, several facets in the implementation of Cuttlefish 3 turned out to provide particularly useful insights. Fast parsing [41] of massive amounts of (compressed) sequences is crucial when dealing with terabytes of data. When partitioning the input sequences to a number of subgraph buckets, these buckets can grow very large and surpass the input sizes, and as such (de)compressing the buckets on-the-fly is important [14]. For in-memory representation of the subgraphs, more cache-friendly data structures have the potential to yield non-trivial performance improvements. Additionally, disk speeds have substantial impact on this I/O heavy paradigm, since disks are frequently used to batch-in and -out partitions of various data structures throughout. We suspect that fast disks, that admit parallel access, will become increasingly important to enable high-performance bioinformatics applications in the future. Finally, while further improvements, of both an engineering and algorithmic nature are certainly possible, and we are continuing to explore these, we have reason to believe that current best-in-class approaches for (colored) compacted de Bruijn graph construction are not far from a reasonable, approximate, practical lower bound on construction time.

## Declarations

R.P. is a co-founder of Ocean Genomics Inc.

## Acknowledgments

This work was supported by the US National Institutes of Health R01HG009937, NSF OAC-2517201, and 2513656, NSF SaTC-2317194, and by grants 2022-311195 and 2024-342821 from the Chan Zuckerberg Initiative DAF, an advised fund of the Chan Zuckerberg Initiative Foundation.

## Footnotes

3 Note that since Cuttlefish 2 does not support color information, we exclude it from our baseline comparisons when evaluating colored compacted de Bruijn graphs.

5 To bound the maximum lengths of super k-mers, we slightly abuse notation such that mm(x) also includes the offset of the minimizer in x, with ties broken in favor of the leftmost offset, and thus µ(x) also includes the offset.

6 The maximum degree of a side of side of vertex is |Σ| =4. Thus each vertex has a maximum degree of 2×|Σ| =8.

7 Some unique identifier of the maximal path p.

8 The ϕ vertex has rank 0 in all paths. The last vertex in a path having rank 0 forms a corner case in the rank computation of edges from its endpoints’ ranks, and a special case with paths having exactly two edges.

9 This statement is not quite true for vertices in 𝒫_i_ forming sub-paths within paths. However, for one endpoint of the sub-paths, the path ID and the rank have been computed at this point, and these values are propagated to the rest of the sub-path.

10 Including the first vertex in the walk.

11 The entire set of h^*′*^(j)’s is not required to be available simultaneously to compute h^*′′*^.

12 A vertex being present in the same source s_j_ multiple times presents a fundamental barrier to the online nature of h^*′′*^ and, as a result, to the uniqueness of h(c) from different multiset variants of c—see Sec. 4.3.

13 L and R are conceptual—all computations are directly over M

14 We devised this method to speed up the partitioning phase of Cuttlefish 3, but note that it corresponds essentially to the “Two-Stacks” method described by Groot Koerkamp and Martayan [36], which they note is well-known in the competitive programming community [37,38]

15 commit ID d69f421a9bf95e9df3188c147191c92d36f9952f

16 v2.0.0

17 We adopt `cpulimit` as we noticed that GGCAT, during extended executions, uses up to 175% of the thread resources requested by the user.

