## Supplementary Material for "Faster and Scalable Parallel External-Memory Construction of Colored Compacted de Bruijn Graphs with Cuttlefish 3"

```
Partition-Graph(𝒮, k)
ℓ = Minimizer-Size(k) // Set a suitable minimizer-size
𝒢 = array of 4^ℓ empty graphs
for each s ∈ 𝒮
    parallel for each (k + 1)-mer e in s
        let e : u ⊙^(k−1) v
        𝒢_rmm(u) = 𝒢_rmm(u) ∪ {e}                    // rmm(u) == lmm(v)
return 𝒢
```

```
Contract-Subgraphs(𝒢, f_0)
// Representation of the discontinuity graph Γ
E = (v + 1) × (v + 1) matrix of empty edge-lists
parallel for each G_i ∈ 𝒢
    M = empty hashtable for (vertex, state) pairs
    for each e ∈ G_i
        let e : u ⊙^(k−1) v
        M_û = δ(M_û, v), M_v̂ = δ(M_v̂, u)

    for each v ∈ Keys(M)
        if not Visited(M_v) and not Isolated(M_v)
            u = Extract-Maximal-Unitig(v)
            let u = (l, . . . , r)
            x = l if lmm(l) ≠ i, ϕ o/w
            y = r if rmm(r) ≠ i, ϕ o/w
            if x == ϕ and y == ϕ
                Output(u)
            else
                e = ({x, y}, 1, Spell(u))
                p = 𝔓(x), q = 𝔓(y)
                E_p,q = E_p,q ∪ {e}                   // WLOG, p ≤ q
return E
```

```
Contract-Paths(Γ)
Π = set of (ν + 1) empty lists of metavertices
for j = ν downto 1
    M = empty hashtable for (vertex, neighbor) pairs
    Δ = Contract-Block(Γ_{j,j}, M)
    parallel for e ∈ Γ_{i,j ∀i<j}
        let e = ({x, y}, w_xy, _)                    // 𝔓(x) = i, 𝔓(y) = j
        if y ∉ Keys(M)
            M_y = (x, w_xy)
        else let M_y : (z, w_yz)
            if x == φ and z == φ
                Add Metavertex(y, w_xy, w_yz) to Π_j
            else p = 𝔓(x), q = 𝔓(z)
                e′ = ({x, z}, w_xy + w_yz)
                Γ_{p,q} = Γ_{p,q} ∪ {e}              // WLOG, p ≤ q
    parallel for each e ∈ Δ
        Contract-Subpath(e) and add any resultant
        meta-vertex to Π
return Π

Expand-Paths(Γ, Π)
Λ = set of (ν + 1) empty lists of edge path-info
for i = 1 to ν
    M = Populate-Hashtable(Π_i)
    Expand-Block(Γ_{i,i}, M)
    parallel for each e ∈ Γ_{i,j ∀j>i}
        let e : ({x, y}, w_xy, s)                    // i == 𝔓(x), j == 𝔓(y)
        let π_x = M_x : (p_x, r_x)
        p_y = p_x
        r_y = (r_x + w_xy) or (r_x − w_xy)           // based on sides
        π_y = (p_y, r_y)
        Add (y, π_y) to P_j
        if w_xy == 1
            λ = Edge-Path-Info(π_x, π_y)
            Λ_i = Λ_i ∪ {(λ, s)}
return Λ

Collate-Unitigs(Λ)
KV = η-sized array of empty lists
parallel for i = 1 to ν
    for each (λ, s) ∈ Λ_i
        let λ : (p, r)
        KV_{h(p)} = KV_{h(p)} ∪ {(p, r, s)}
𝒫 = Φ
parallel for i = 1 to η
    let KV_i : {(p, r, s)}
    Sort(KV_i) on (p, r)
    for each maximal stretch KV_[a,b] with same p
        𝒫 = 𝒫 ∪ {s_a ⊙^{k−1} ... ⊙^{k−1} s_b}
return 𝒫
```

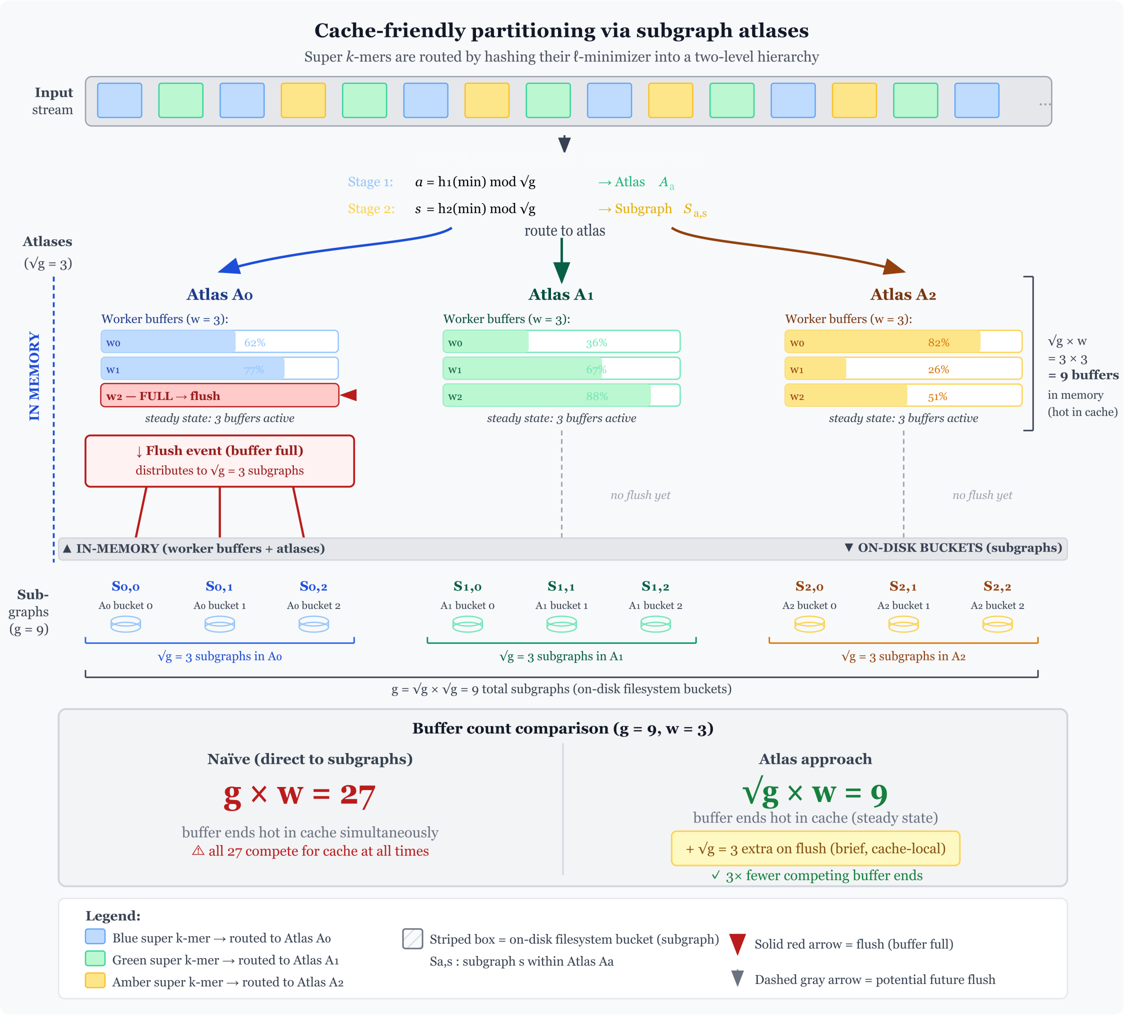


Fig. S1: An overview illustrating the subgraph atlas approach for cache-friendly super k-mer partitioning. The approach minimizes the number of buffers that must be retained in memory to collate output before it is flushed to the appropriate subgraph file (bucket). By minimizing the number of distinct buffers, we improve caching behavior while still minimizing lock contention.

### 0.1 Proofs

We slightly abuse the notations here as, based on the context, minimizer instances $m$'s to also include their positions.

**Lemma 1.** *Only the last and the first* $(k-1)$*-mers in a super* $k$*-mer can have minimizer instances different to that of the super* $k$*-mer.*

*Proof.* The claim trivially holds true for super $k$-mers of length $k$. Let a super $k$-mer of length $> k$ be $s = v_0 \odot^{k-1} \ldots \odot^{k-1} v_n$ with minimizer instance $m$. So $\forall_i \mu(v_i) = m$. Let us express $s$ in terms of $(k-1)$-mers as $s = u_0 \odot^{k-2} \ldots \odot^{k-2} u_{n+1}$, where the $u_i$'s are $(k-1)$-mers.

Suppose that for some $0 < j < n+1$, $\mu(u_j) = m' \neq m$. Note that $u_j = \mathtt{pre}_{k-1}(v_j)$, and hence $\mu(\mathtt{pre}_{k-1}(v_j)) = m' \neq m$. Since $\mu(v_j) = m$, two cases are possible for the position of $m$:

1. $m$ is the last $\ell$-mer of $v_j$. This implies that $m$ is absent in $v_{j-1}$. So $\mu(v_{j-1}) \neq m$, a contradiction.
2. Or, $m$ is present in $\mathtt{pre}_{k-1}(v_j)$. As we know $\mu(\mathtt{pre}_{k-1}(v_j)) = m'$, it must be that $m' < m$. This implies that there exists an $\ell$-mer $m' < m$ in $v_j$. Hence $\mu(v_j) \neq m$, another contradiction.

Thus, for $0 < j < n+1$, no $(k-1)$-mer $u_j$ has a minimizer $m' \neq m$.

**Corollary 1.** *All the edges induced from a super* $k$*-mer belong to the same subgraph.*

*Proof.* Suppose that for a super $k$-mer $s = v_0 \odot^{k-1} \ldots \odot^{k-1} v_n$, its minimizer instance is $m$, and the edge $e = v_i \odot^{k-1} v_{i+1}$ belongs to some $\mathcal{G}_{m'} \neq \mathcal{G}_m$. This implies $\mathtt{rmm}(v_i) = \mathtt{lmm}(v_{i+1}) = m' \neq m$. This is impossible per lemma 1. Hence $e$ belongs to $\mathcal{G}_m$.

**Lemma 2.** *A discontinuity vertex has edges incident to it at one side only at some subgraph* $\mathcal{G}_i$*.*

*Proof.* For a discontinuity vertex $v$, suppose the edge $e = u \odot^{k-1} v$ exists in the input. Then $v$ exists in the subgraph $\mathcal{G}_{\mathtt{lmm}(v)}$. WLOG, let $e$ be incident to the $\mathtt{front}$ of $v$, i.e. $v = \widehat{v}$.

Now assume that another edge $e'$ exists at the $\mathtt{back}$ of $v$ within $\mathcal{G}_{\mathtt{lmm}(v)}$. Hence $e'$ has the form $e' = \widehat{v} \odot^{k-1} u'$. So $e'$ belongs to $\mathcal{G}_{\mathtt{rmm}(\widehat{v})}$. This implies that $\mathcal{G}_{\mathtt{lmm}(\widehat{v})} = \mathcal{G}_{\mathtt{rmm}(\widehat{v})}$, i.e. $\mathtt{lmm}(\widehat{v}) = \mathtt{rmm}(\widehat{v})$. Thus $v$ is not a discontinuity vertex—a contradiction. So $v$ cannot have edges at both sides in some $\mathcal{G}_i$.

**Lemma 3.** *A vertex in* $\Gamma$ *has at most one edge at a side.*

*Proof.* Suppose not for some vertex $v$. Let $v$ have two edges $e_1$ and $e_2$ in $\Gamma$ at side $s_v$. $e_1$ corresponds to a locally maximal unitig $p_1$ at a subgraph $\mathcal{G}_{m_1}$ and $e_2$ to $p_2$ at $\mathcal{G}_{m_2}$. $m_1 \neq m_2$, because otherwise the same subgraph introduces both $e_1$ and $e_2$, and $v$ cannot belong to two maximal unitigs in the same subgraph.

WLOG, assume $s_v = \mathtt{front}$. So $p_1$ and $p_2$ both connect to the $\mathtt{front}$ of $\widehat{v}$. Since $\widehat{v}$ is a discontinuity vertex, it can have edges only incident at its $\mathtt{front}$ at $\mathcal{G}_{\mathtt{lmm}(\widehat{v})}$ and only at its $\mathtt{back}$ at $\mathcal{G}_{\mathtt{rmm}(\widehat{v})}$ (see Lemma 2). This implies that $p_1$ and $p_2$ both belong to the same subgraph $\mathcal{G}_{\mathtt{lmm}(\widehat{v})}$. Thus $m_1 = m_2$, which is a contradiction. So $v$ has at most one edge at $s_v$ in $\Gamma$.

**Lemma 4.** $\Gamma$ *is a collection of disjoint maximal paths only touching at the sentinel vertex* $\phi$*.*

*Proof.* Suppose not. Then there exists two maximal paths $p_1$ and $p_2$ connected with an edge $e = \{u, v\}$. Let $u \in p_1$ and $v \in p_2$. Let $e$ be incident to the side $s_u$ of $u$. Then $s_u$ must have an incident edge $e' \neq e$ in $p_1$, because otherwise $p_1$ could have been extended through $e$ and joined with $p_2$. This implies that $u$ has $> 1$ edges incident at $s_u$. This is a contradiction per Lemma 3. So $p_1$ and $p_2$ are disjoint.

**Lemma 5.** *A maximal unitig is* $G$ *with at least one discontinuity* $k$*-mer corresponds to a maximal path* $p = (\phi, v_1, v_2, \ldots, v_n, \phi)$ *in* $\Gamma$*.*

*Proof.* Let $u$ be a maximal unitig in $G$, and its decomposition into maximal super $k$-mers be $u = (u_1 \odot^{k-1} u_2 \odot^{k-1} \ldots \odot^{k-1} u_n)$, with $u_i$ having minimizer $m_i$. Consider the rightmost $k$-mer $r$ of $u_i$ and the leftmost $k$-mer $l$ of $u_{i+1}$. Either one of $l$ and $r$ is a discontinuity $k$-mer, because otherwise $l$ and $r$ have the same minimizer and thus $u_i$ and $u_{i+1}$ are not maximal. That vertex $y$ is present in $\mathcal{G}_{m_i}$ (and in $\mathcal{G}_{m_{i+1}}$). Similarly for $u_{i-1}$ and $u_i$, there is a discontinuity vertex $x$ at their border, and $x$ is present in $\mathcal{G}_{m_i}$ (and in $\mathcal{G}_{m_{i-1}}$). The sequence between $x$ and $y$, inclusive, is constructed as a maximal unitig in $\mathcal{G}_{m_i}$, and the edge $\{x, y\}$ is added to $\Gamma$. These edges produced from the $u_i$'s form a maximal path in $\Gamma$.

### 0.2 Colored Compacted de Bruijn Graph Construction

We evaluate CUTTLEFISH 3 compared to GGCAT with $k = 31$ to construct the colored de Bruijn graph on the genome sequence datasets. Table S1 contains the benchmark results.

Table S1: Time (in wall clock), memory (in GB, in parentheses left to the vertical bar), and intermediate disk-usage (in GB, in parentheses right to the vertical bar) performance results for constructing colored compacted de Bruijn graphs with 32 threads. The best metrics in each row are highlighted.

| k | Dataset | GGCAT | Cuttlefish 3 |
|---|---|---|---|
| 31 | Human gut | 19 m (**20.6** \| 77) | **11 m** (25.2 \| 102) |
| | Salmonella (150K) | 02 h 18 m (10.4 \| 179) | **42 m** (**10.3** \| 511) |
| | Salmonella (309K) | 05 h 18 m (**14.0** \| 374) | **01 h 32 m** (15.9 \| 1054) |
| | Bacterial archive (661K) | 13 h 29 m (35.8 \| 1300) | **03 h 18 m** (**33.7** \| 2190) |
| 63 | Human gut | 15 m (22.2 \| 62) | **11 m** (**20.4** \| 74) |
| | Salmonella (150K) | 02 h 21 m (**10.1** \| 127) | **39 m** (11 \| 380) |
| | Salmonella (309K) | 05 h 39 m (**14.2** \| 257) | **01 h 30 m** (17.5 \| 783) |
| | Bacterial archive (661K) | 11 h 25 m (**36.2** \| 875) | **03 h 15 m** (41.3 \| 1602) |

Table S2: Timing (in wall clock) performance results for constructing colored compacted de Bruijn graphs across different numbers of threads.

| Threads | GGCAT | Cuttlefish 3 |
|---|---|---|
| 1 | 3698 | 4942 |
| 2 | 2345 | 2998 |
| 4 | 1080 | 1506 |
| 8 | 573 | 829 |
| 16 | 311 | 464 |
| 32 | 177 | 273 |

### 0.3 Parallel Scaling

### 0.4 Tools and Execution Commands

For the experiments, we used the following versions of the tools: GGCAT (v2.0.0) and Cuttlefish 3 (commit ID d69f421). We adopt `cpulimit` to restrict the CPU-usage of the tools to $t + 1$ threads for a request of t threads [3] as we noticed that GGCAT, during extended executions, uses up to 175% of the thread resources requested by the user.

The following commands have been used in executing the tools.

1. GGCAT: `cpulimit -f -l ${cpu} -z -- ggcat build -l ${input_list} -k ${k} -s 1 -j ${threads} -t ${temp_dir} -o ${output_file} -c`
2. Cuttlefish 3: `PARLAY_NUM_THREADS=${threads} cpulimit -f -l ${cpu} -z -- cuttlefish build -l ${input_list} -k ${k} -w ${temp_dir} -o ${output_file} --ref --color`

[3] It is customary to use an extra thread for various background read/write operations.
